## Supplementary information for "Structural basis for two-way communication between dynein and microtubules"

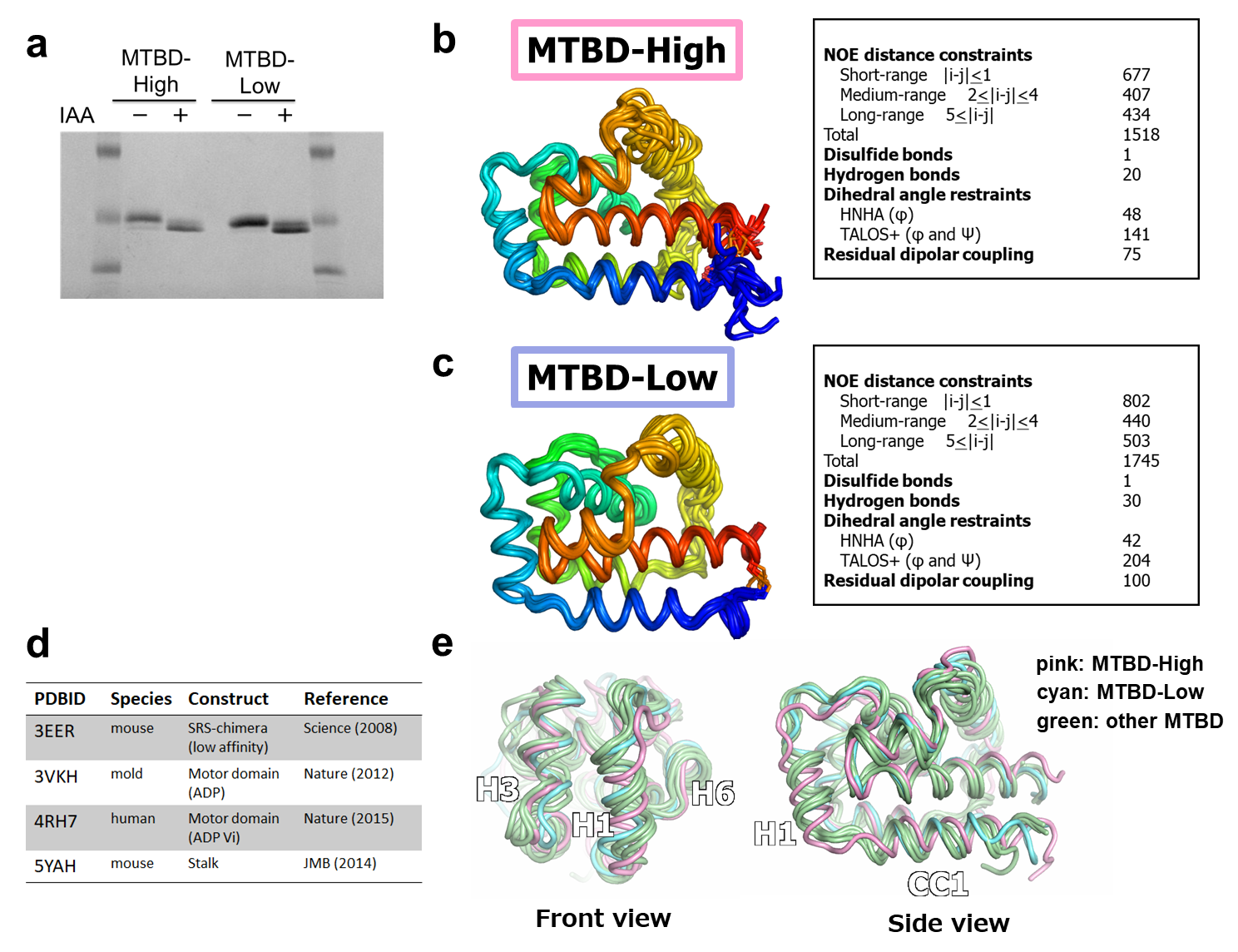

**Supplementary Figure 1.**

**The NMR structures of MTBD-High and MTBD-Low**

(a) SDS-PAGE gel of MTBD-High and MTBD-Low. The samples were resuspended in the standard SDS loading buffer in the presence (+) and absence (-) of 5 mM iodeactoamide (IAA). (b, c) NMR structures of MTBD-High and MTBD-Low. Backbone trace of the ten lowest-energy structures and the distance constraints used for the Xplor-NIH calculation of the MTBD-High (b) and MTBD-Low (c) structures. (d, e) Comparison of the NMR structures of MTBD-High and MTBD-Low with previously reported crystal structures of MTBD. (d) A table summarizing the previously determined crystal structures of dyneins containing the MTBD moiety^1–4^. (e) Superposition of MTBD-Low (cyan), MTBD-High (pink), and the MTBD moiety of the previous crystal structures (light green).

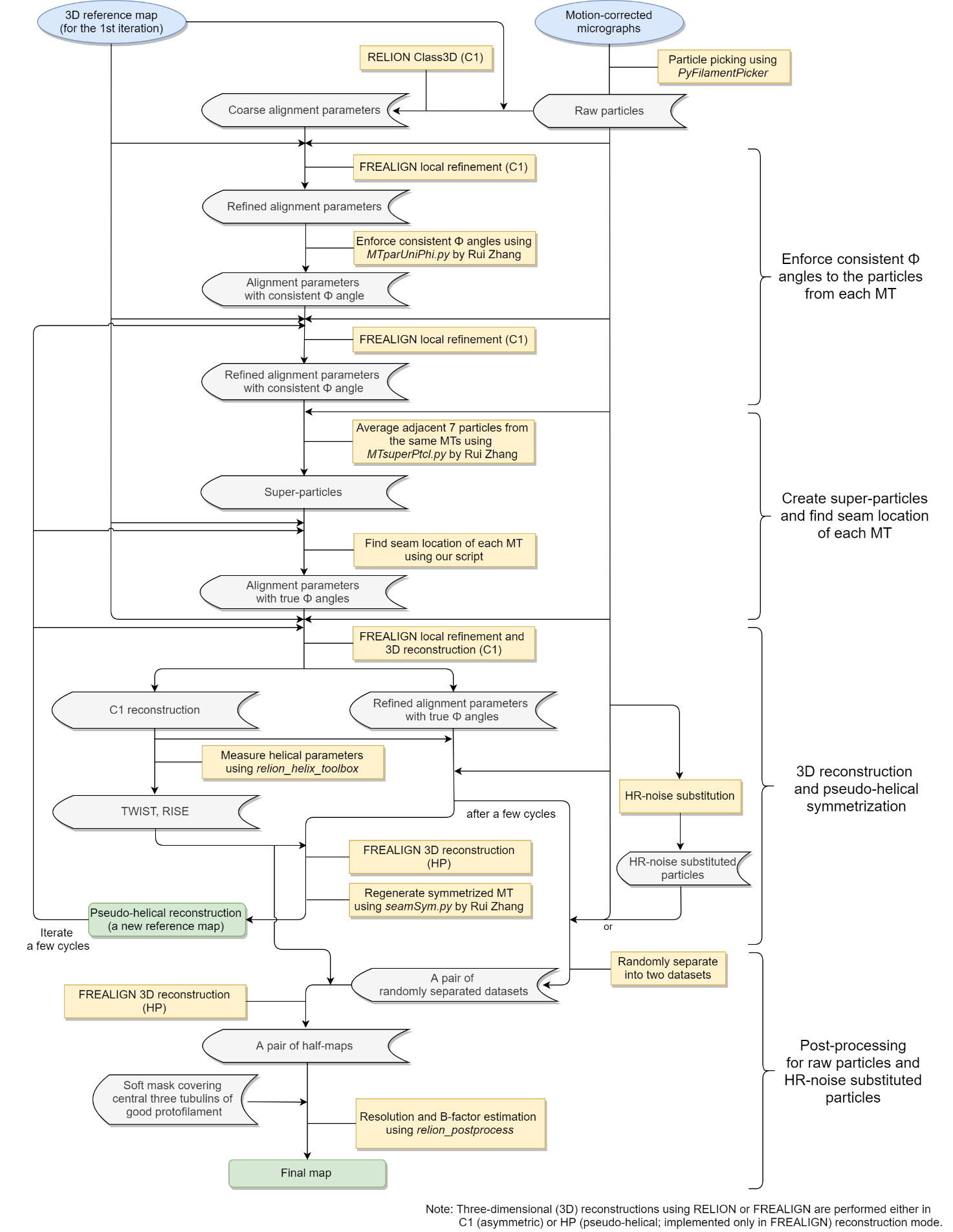

**Supplementary Figure 2**.

### **Processing workflow for the cryo-EM structures of MTBD-High-MT complexes**

Seam determination was performed using the “super-particle”-based approach with a few modifications using original scripts developed in our lab. In the post-processing steps, Fourier Shell Correlation (FSC) calculation and B-factor estimation were performed using *relion_postprocess* with a soft mask covering the central three tubulins of one “good” protofilament. To calculate FSC_true_, we calculated both FSC_t_ from the raw dataset and FSC_n_ from the high-resolution (HR) noise-substituted dataset.

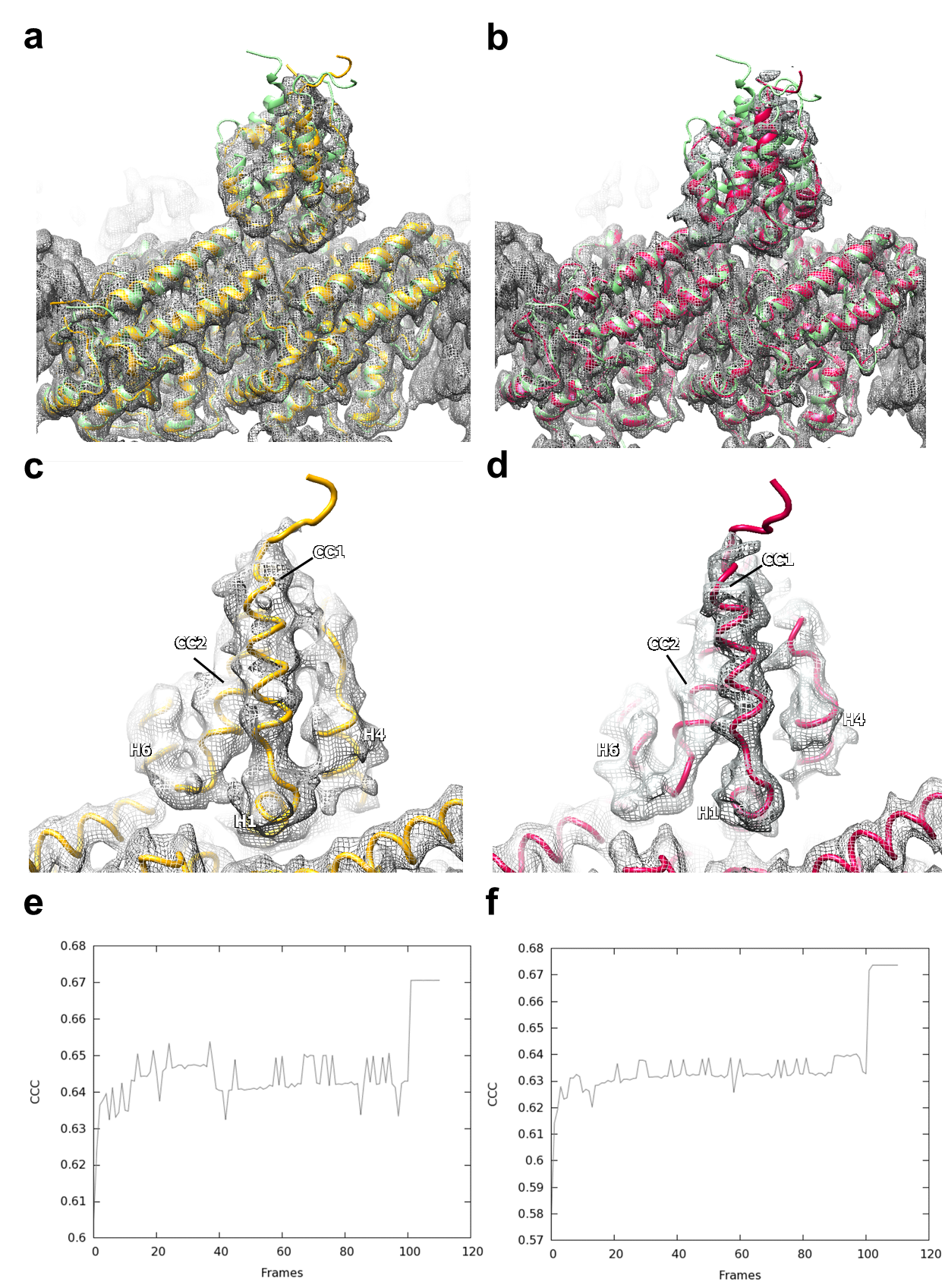

**Supplementary Figure 3**

**Flexible fitting of MTBD-High and tubulin dimer into the cryo-EM maps of MTBD-High-MT complex in the absence (-) and presence (+) of DTT.**

(a, b) The molecular dynamics flexible fitting (MDFF) of MTBD-High and the tubulin dimer (PDB code: 1JFF) into the cryo-EM maps filtered to 4 Å: (a) for DTT(-) and (b) for DTT(+).The initial placement of the complex is shown in light green. (c, d) Close-up views of the CC1 moiety fit into the cryo-EM maps filtered to 4 Å: (c) for DTT(-) and (d) for DTT(+).

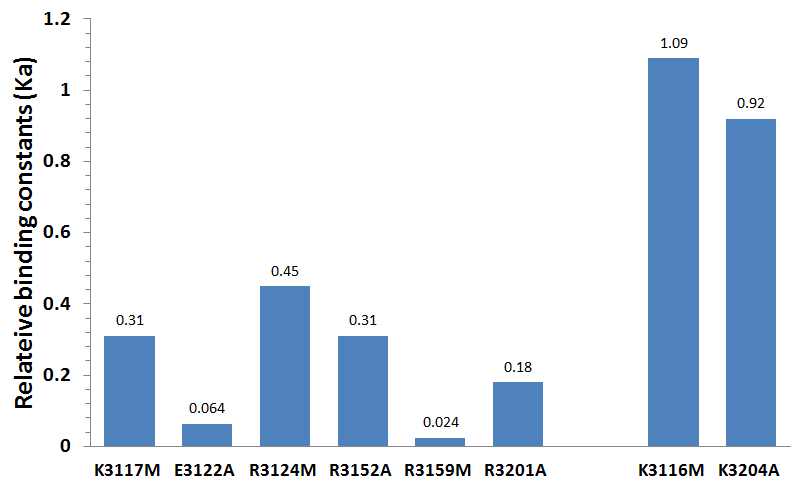

**Supplementary Figure 4.**

**Mutational analysis of MTBD-High.**

The binding constants (normalized to those of MTBD-High) for each mutant are plotted as a bar graph. We confirmed that the overall conformation of MTBD was not disrupted by the mutation based on the HSQC (heteronuclear single quantum coherence) spectra of mutants.

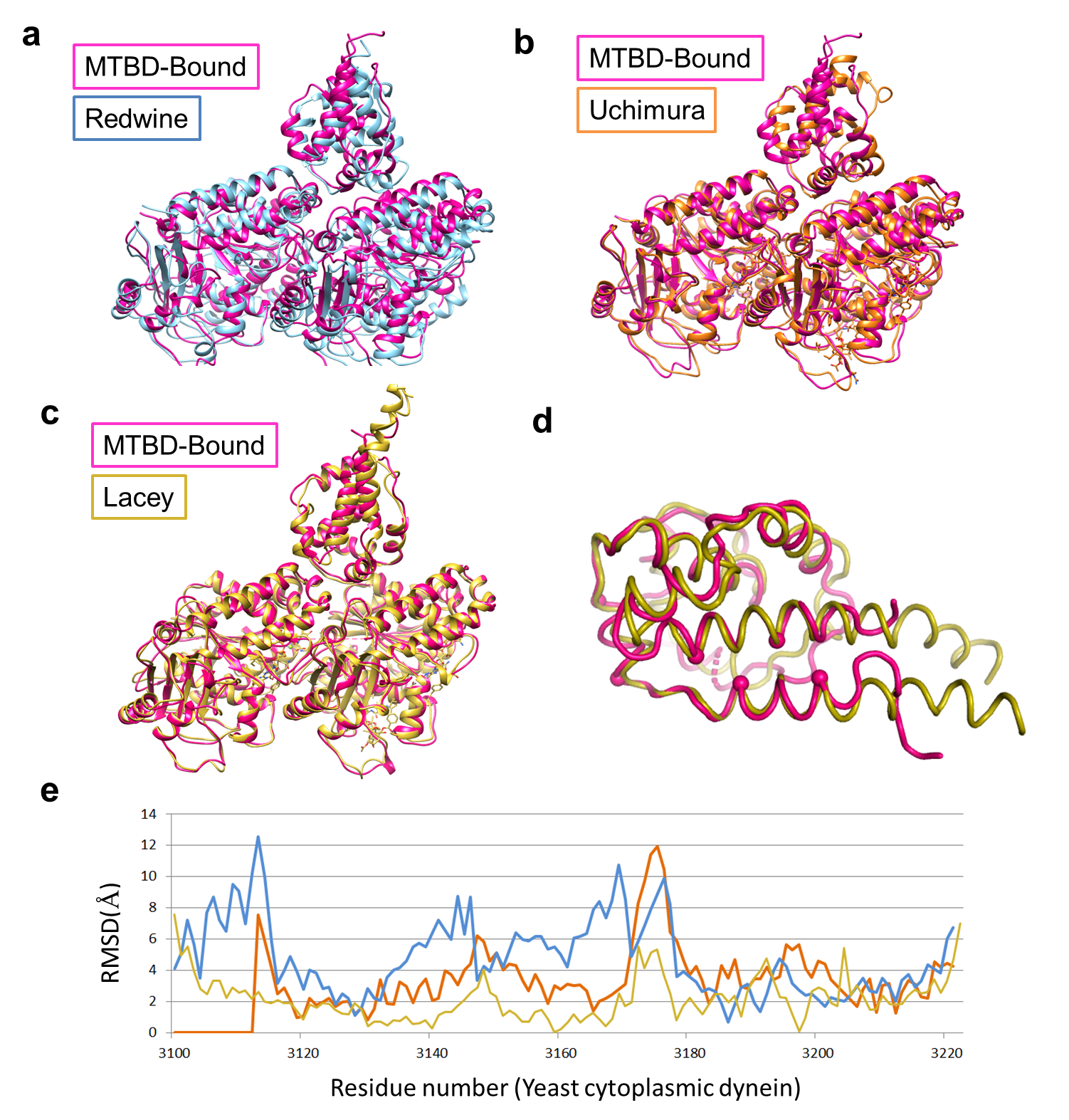

**Supplementary Figure 5.**

**Comparison of overall structure of three cryo-EM-based models of dynein MTBD-MT complexes.**

(a-c) Superimposition of the cryo-EM based models of (a) Redwine et al.^5^ (cyan), (b) Uchimura et al.^6^ (orange), and (c) Lacey et al.^7^ (yellow) with the model derived in the current study (magenta). All structures are aligned with respect to the α-tubulin subunit. (d) Superposition of MTBD moiety between the Lacey model (yellow) and that of the present study (magenta). (e) Plots of RMSD values for the Cα position of each MTBD residue in the current model and in those of the Redwine (cyan), Uchimura (orange), and Lacey (yellow) models.

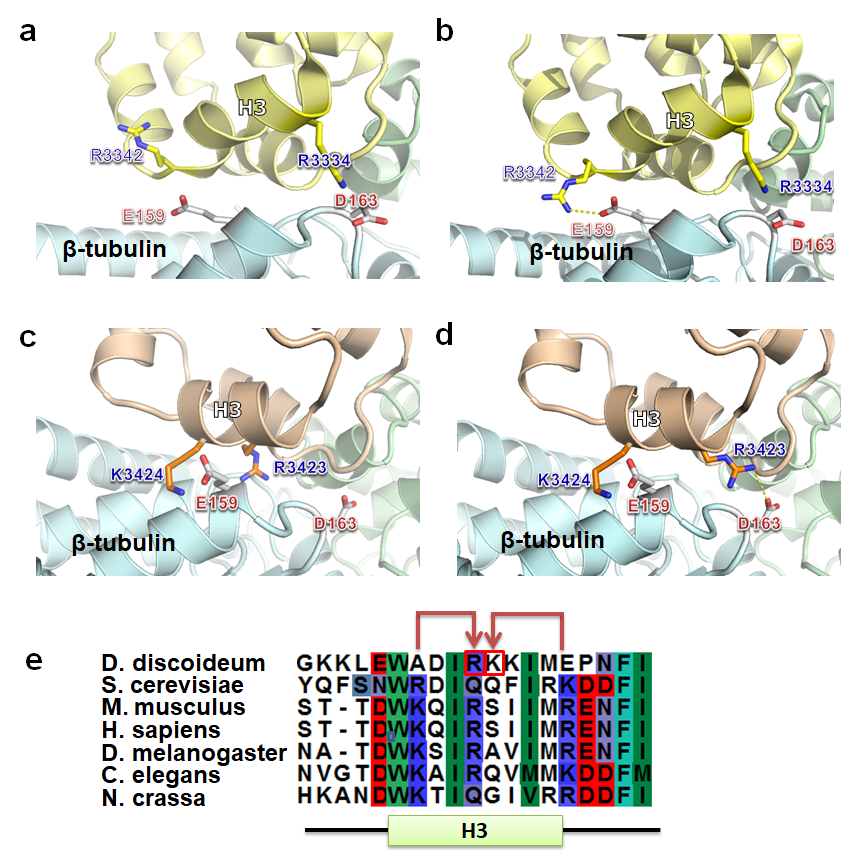

**Supplementary Figure 6.**

**Conservation of the interaction mode of H3**

(a) The conserved H3 residues forming salt bridges of mouse cytoplasmic dynein are shown by stick models. (b) The manipulated side chain of R3342 re-orients toward E159, which is sufficiently close to form a salt bridge. (c) The conserved H3 residues forming salt bridges of Dictyosterium cytoplasmic dynein are shown by stick models. (d) The manipulated side chain of R3423 re-orients toward D163, which is sufficiently close to form a salt bridge. (e) Sequence alignments of the H3 region of the cytoplasmic dyneins of various species.

### **Supplementary Table 1.**

### **Statistics of the cryo-EM data collection and image analysis for MTBD-High-MT complex in the absence (-) and presence (+) of DTT**

|  | | MTBD-MT DTT(-) | MTBD-MT DTT(+) |
| --- | --- | --- | --- |
| Number of micrographs | | 1,820 | 621 |
| Number of MTs | Initial | 2,479 | 1,427 |
|  | Final | 1,920 | 1,044 |
| Number of particles | Initial | 76,377 | 35,636 |
|  | Final | 58,999 | 32,666 |

### **Supplementary Table 2.**

### **Fourier Shell Correlations (FSC) and map sharpening B-factor for the 3D reconstructions of MTBD-High-MT complex in the absence (-) and presence (+) of DTT**

|  | | MTBD-MT DTT(-) | MTBD-MT DTT(+) |
| --- | --- | --- | --- |
| Resolution (Å)  (asymmetric reconstruction, C1) | FSC_true_ | 4.44 | 4.33 |
|  | FSC_t_ | 4.47 | 4.29 |
|  | FSC_n_ | 8.80 | 9.10 |
| Resolution (Å)  (pseudo-helical reconstruction, HP) | FSC_true_ | 3.94 | 3.66 |
|  | FSC_t_ | 3.94 | 3.62 |
|  | FSC_n_ | 7.76 | 8.00 |
| Map sharpening  B-factor (Å^2^) | C1 | -91.3 | -82.0 |
|  | HP | -145.0 | -128.9 |

### **Supplementary Video 1.**

### **Continuous movements of CC1 and H1 from MTBD-Low to MTBD-High to MTBD-Bound.**

A morphed series showing the conformational transition of the MTBD from MTBD-Low to MTBD-High to MTBD-Bound. Orientation and coloring are the same as those used in Figure 2d.
